# Transport-Related Effects on Intrinsic and Synaptic Properties of Human Cortical Neurons: A Comparative Study

**DOI:** 10.1101/2024.10.30.621044

**Authors:** Guanxiao Qi, Danqing Yang, Aniella Bak, Werner Hucko, Daniel Delev, Hussam Hamou, Dirk Feldmeyer, Henner Koch

**Affiliations:** Institute of Neuroscience and Medicine, INM-10, Research Centre Juelich, Jülich, Germany; Department of Epileptology and Neurology, RWTH Aachen Uniklinik, Aachen, Germany; Department of Neurosurgery, RWTH Aachen Uniklinik, Aachen, Germany; Department of Psychiatry, Psychotherapy and Psychosomatics, RWTH Aachen University, Aachen, Germany; Jülich-Aachen Research Alliance-Brain, Translational Brain Medicine, Aachen, Germany; Department of Neurosurgery, University Hospital Erlangen, Friedrich-Alexander University, Erlangen Nürnberg, Erlangen, Germany

## Abstract

Transporting human brain tissue from the operating theater to an off-site laboratory may affect sample integrity for electrophysiological studies. This study investigated how a 30-40 minute transport influenced the intrinsic, synaptic, and morphological properties of human cortical neurons. Electrophysiological recordings were performed on Layer 2/3 (L2/3 pyramidal cells and fast-spiking (FS) interneurons from human cortical slices (n = 200 neurons from 32 surgeries), comparing on-site recordings at RWTH Aachen University Hospital and off-site at Research Centre Juelich. Action potential firing patterns remained largely preserved across both recording sites, but several differences were observed. Off-site recorded pyramidal cells showed a slightly depolarized resting membrane potential and a significantly lower rheobase current. In off-site recorded FS interneurons, we found a narrower action potential half-width and an increased amplitude, suggesting altered ion channel kinetics and/or neuromodulatory environment. Additionally, a significant reduction in large rhythmic depolarizations (LRDs) and the amplitudes of excitatory postsynaptic potentials (EPSPs) in off-site recorded FS interneurons indicated an impaired synaptic efficacy. The dendritic spine densities in apical oblique and apical tuft dendrites of off-site recorded pyramidal cells were also reduced. These findings emphasize the need for optimized transport conditions to preserve synaptic integrity, network properties, and neuronal morphology. Standardized protocols are crucial for ensuring reliable and reproducible results in studies of human cortical microcircuits.

**Significance Statement:** This study demonstrates that transporting live human brain tissue for neuronal recordings significantly impacts the intrinsic, synaptic, and network properties of cortical neurons. By comparing on-site and off-site recordings, we found that even a brief transportation (30-40 minutes) induces increased neuronal excitability, reduced synaptic efficacy, and diminished network events such as LRDs. These alterations are likely due to the mechanical stress and washout of critical neuromodulators, which compromise tissue integrity and neuronal function. The findings underscore the necessity for optimizing transport protocols to preserve synaptic and network integrity, ensuring reliable and reproducible results in human brain research. Ultimately, this work advances our understanding of cortical microcircuitry and informs best practices for handling human brain tissue in experimental settings.

## Introduction

The investigation of human cortical microcircuits is attracting increasing interests within the neuroscience community, already spanning a broad range of studies focusing on cellular functions (Peng et al., 2019;Verhoog et al. 2013; Gidon et al. 2020), distinct cell types (Hodge et al., 2019; Yuste et al., 2020; Lee et al., 2023), the occurrence and modulation of network activity (Köhling et al., 1998; Yang et al., 2024), and the underlying mechanisms of both physiological and pathological conditions (Köhling et al., 1999; Gorji et al., 2001; Marcuccilli et al., 2010; Blauwblomme et al., 2019; Barth et al., 2021; Taylor et al., 2024). Given this growing interest, there is a parallel need to optimize procedures for obtaining the highest quantity, best quality, and longest survival of human brain samples, which is crucial for advancing research in this field (Bak et al. 2024; Schwarz et al. 2017, Schwarz et al. 2019, Kraus et al. 2020; Straehle et al. 2023; Wickham et al. 2018).

In *ex vivo* animal experiments, the location of sample collection and the site of experiments can be designed to be in close proximity, thereby largely eliminating the need to optimize sample transportation. In contrast, human tissue samples are obtained in the operating theater of a clinic and consequently, the samples need to be transported from the surgery location to the experimental laboratory. The duration of this transport can vary considerably, from minutes (Bak et al. 2024) to hours (Mittermaier et al. 2024), depending on the location of the experimental setup.

Especially for longer distances, elaborate portable devices for blocks and/or slices of human brain samples were developed and tested before (Köhling et al. 1996; Mittermaier et al. 2024). While these studies showed that brain samples can remain viable for successful electrophysiological recordings from cortical neurons even after long distance transport, more subtle changes may still occur during the transportation of the tissue.

In light of this idea, we aimed to systematically evaluate the impact of the transportation process on the intrinsic and synaptic properties of human cortical neurons. We compared two distinct settings: on-site analysis, where samples are rapidly transferred to a nearby laboratory in the same hospital, and off-site analysis, where samples undergo longer transport duration to a distant laboratory. By performing electrophysiological recordings in both scenarios, we aimed to assess potential differences in cellular properties, synaptic activity, and overall neuronal viability. This comparative analysis provides critical insights into how transportation affects the quality and integrity of human brain samples, guiding future optimization of protocols for corresponding studies.

Our findings reveal that while firing patterns of human cortical cells remain largely unchanged, significant alterations in several cellular parameters occur following the transport to an off-site laboratory. Pyramidal neurons exhibited a more depolarized resting membrane potential and decreased rheobase current along with decreased spine density when recorded off-site, and fast-spiking interneurons displayed an increased action potential amplitude, decreased half-width, and decreased EPSP amplitude. Furthermore, we observed a reduction of spontaneous network events recently described in human slices (Yang et al., 2024), known as large rhythmic depolarizations (LRDs), indicating that synaptic connectivity and activity are particularly susceptible to the effects of transport.

## Methods

### Preparation of acute human slices

Approval (EK-067/20) of the ethics committee of the University of Aachen as well as written informed consent was obtained from all patients, allowing spare, non-pathologic tissue, which was acquired during the surgical approach, to be included in the study. We collected data from the tissue samples of 32 patients (15 females, 17 males; age ranging from 9 to 75 years) (Tab. 1). Tissue preparation was performed according to published protocols (Bak et al. 2024, Yang et al. 2024) (**Fig. 1A)**. In brief, the arachnoid was opened with a scalpel and micro scissors. The edges around the tissue are prepared with a dissector and ultrasonic tissue ablation. Finally, the arterial vessels to the tissue are closed and cut off with micro scissors to shorten the hypoxia time as much as possible and directly transferred into ice-cold ( ∼ 4 ° C) choline-based artificial slicing cerebrospinal fluid (aCSF) (in mM: 110 choline chloride, 26 NaHCO_3_, 10 D-glucose, 11.6 Na-ascorbate, 7 MgCl_2_, 3.1 Na-pyruvate, 2.5 KCl, 1.25 NaH_2_PO_4_, und 0.5 CaCl_2_) equilibrated with carbogen (95% O_2_, 5% CO_2_) and immediately (within 5-10 minutes) transported to the laboratory. The tissue was always kept submerged in cooled and oxygenated slicing aCSF. After removal of the pia, tissue blocks were trimmed perpendicular to the cortical surface and 300 µm thick slices were prepared using a vibratome (Leica VT1200) in either the choline-based slicing aCSF or in ice-cold sucrose-based aCSF containing 206 mM sucrose, 2.5 mM KCl, 1.25 mM NaH_2_PO_4_, 3 mM MgCl_2_, 1 mM CaCl_2_, 25 mM NaHCO_3_, 12 mM N-acetyl-L-cysteine, and 25 mM glucose (325 mOsm/l, pH 7,45). During slicing, the solution was constantly bubbled with carbogen gas (95% O_2_ and 5% CO_2_). Subsequently, slices were first incubated for 30 min at 32-33 °C and then at room temperature in aCSF containing (in mM): 125 NaCl, 2.5 KCl, 1.25 NaH_2_PO_4_, 1 MgCl_2_, 2 CaCl_2_, 25 NaHCO_3_, 25 D-glucose, (300 mOsm/l; 95% O_2_ and 5% CO_2_).

**Fig. 1.**
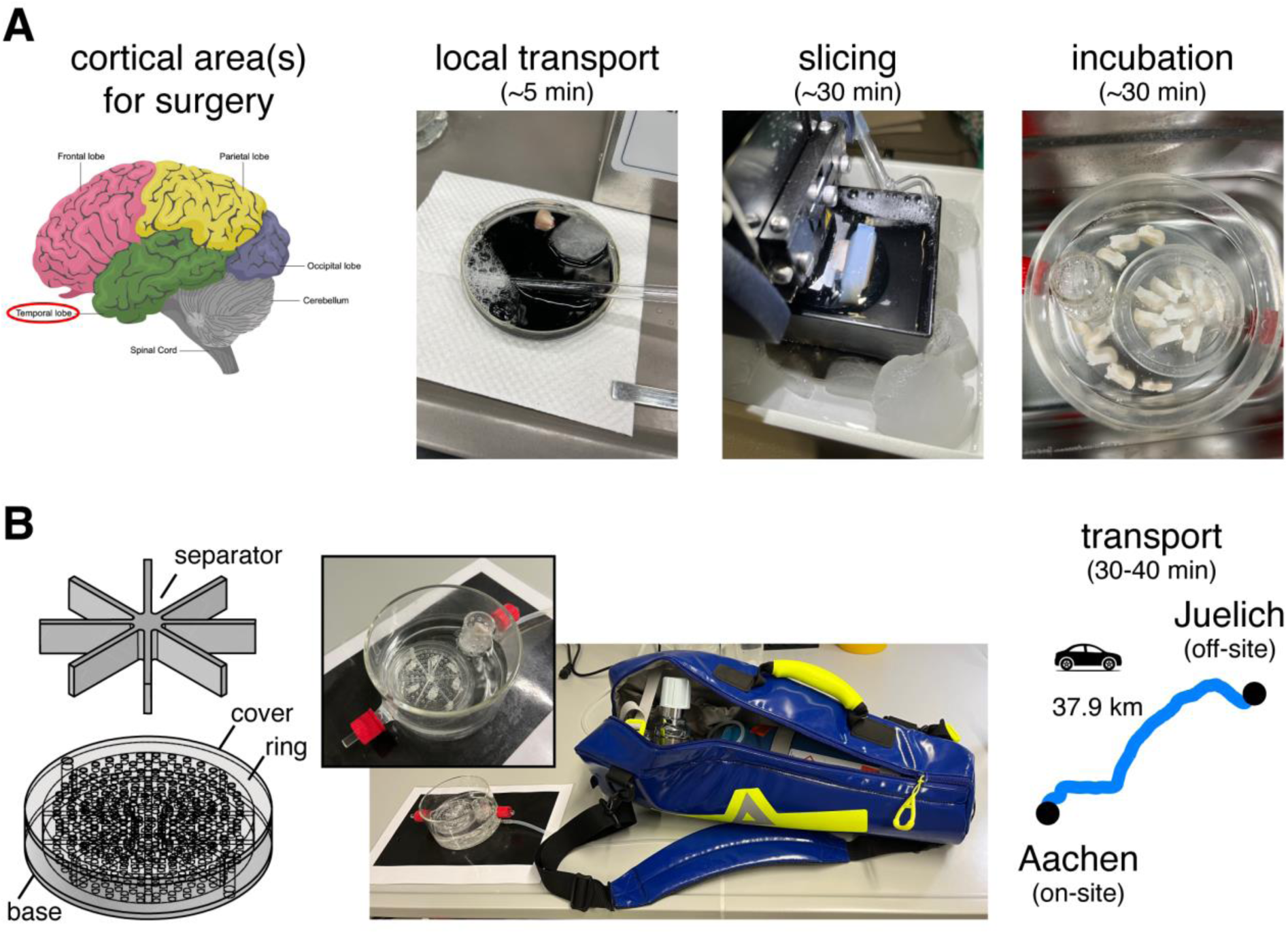
A platform for simultaneous on– and off-site recordings of human cortical neurons ex vivo. (A) Acute human cortical slices were prepared from surgically removed cortical tissue for treatment of epilepsy or brain tumor in the laboratory at RWTH Aachen University Hospital. (B) After slicing and incubation, some slices were kept in the laboratory where they were prepared for recordings. Other slices was transported by car to a laboratory at the Research Centre Juelich for patch-clamp recordings. During transportation, slices were either kept in a custom-made slice keeper (a specialized design of slice holder for transportation is shown on the left) continuously oxygenated with a portable carbogen gas bottle or stored in a 50 ml centrifuge tube filled with oxygen-saturated aCSF.

**Table 1.**
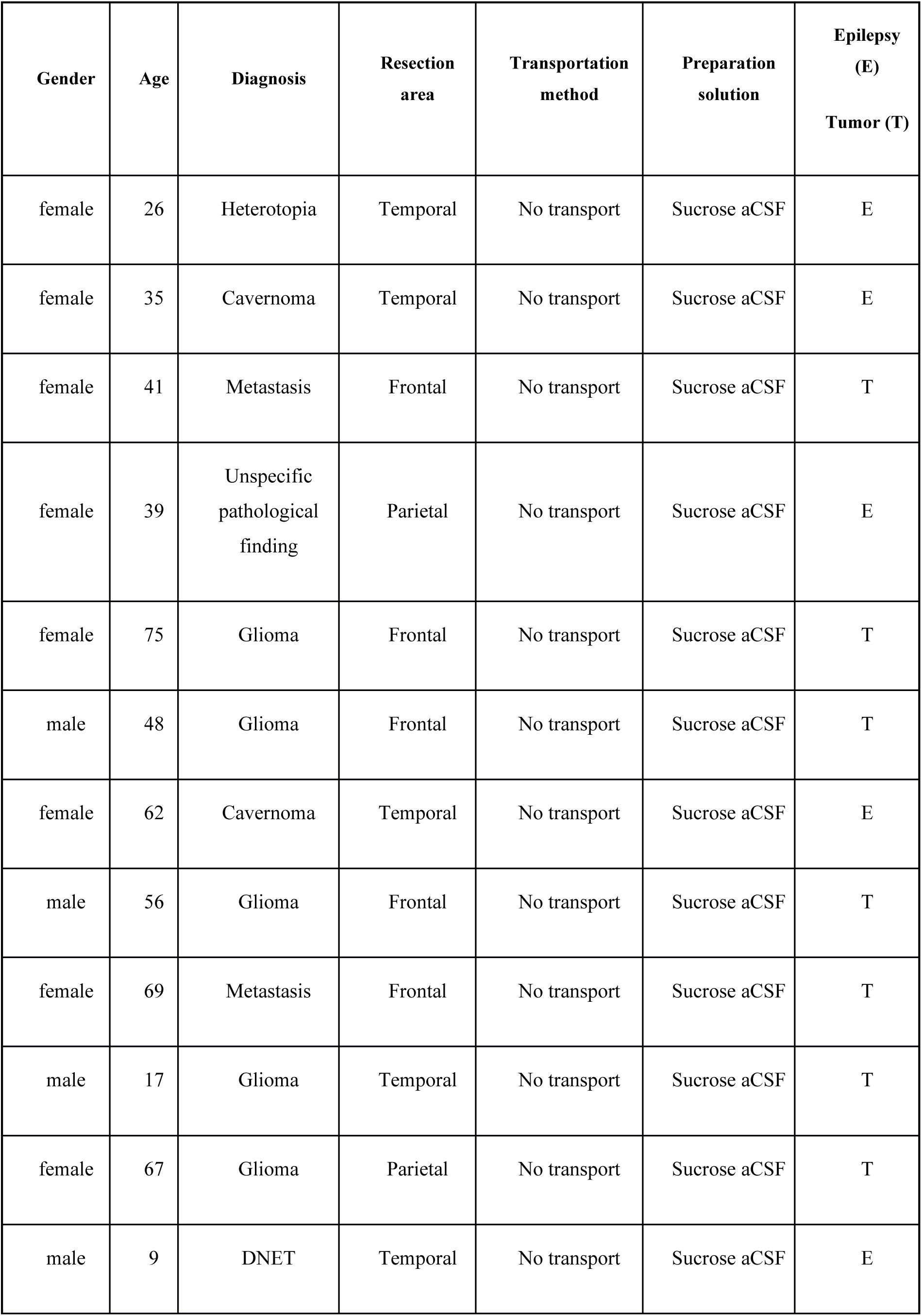
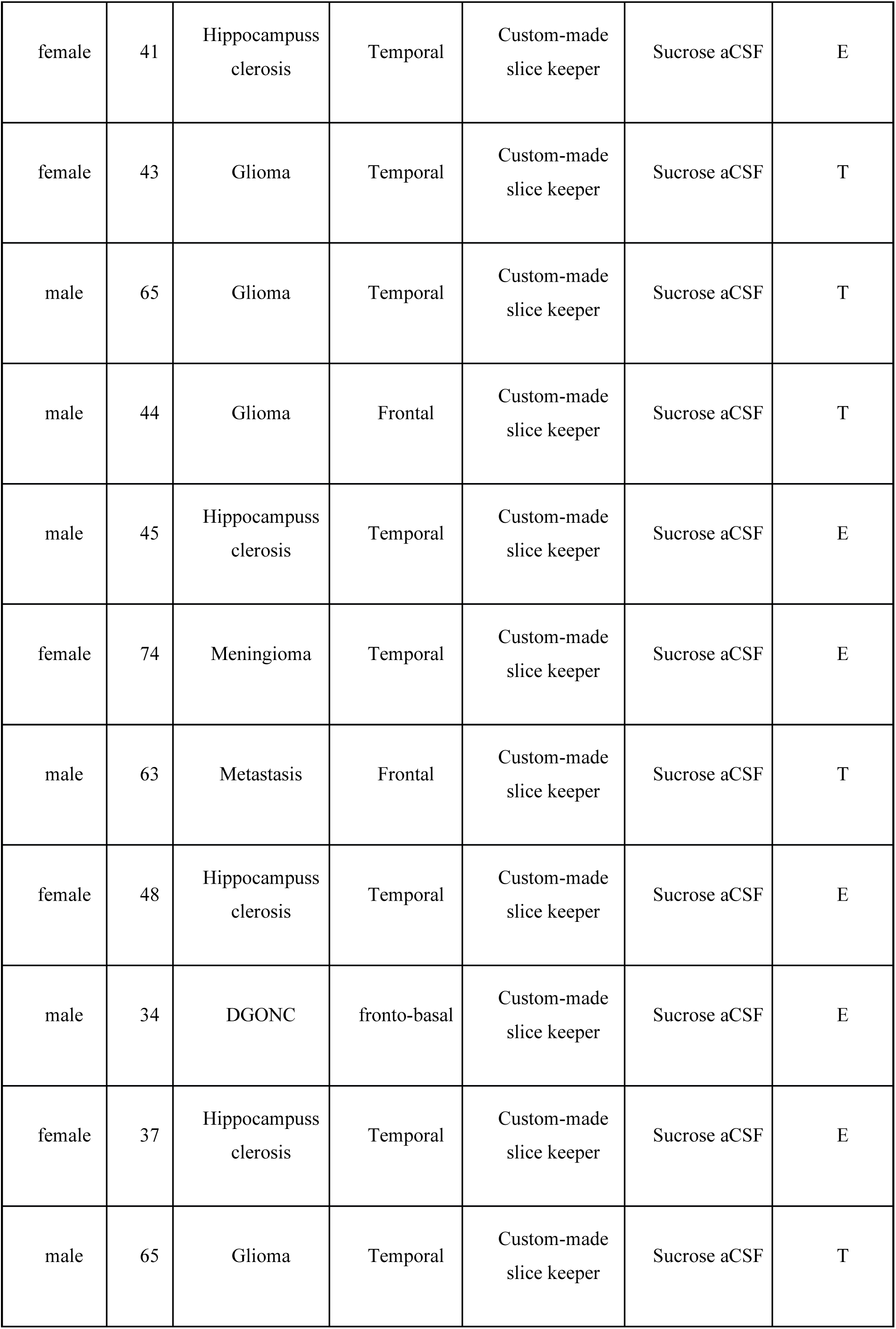
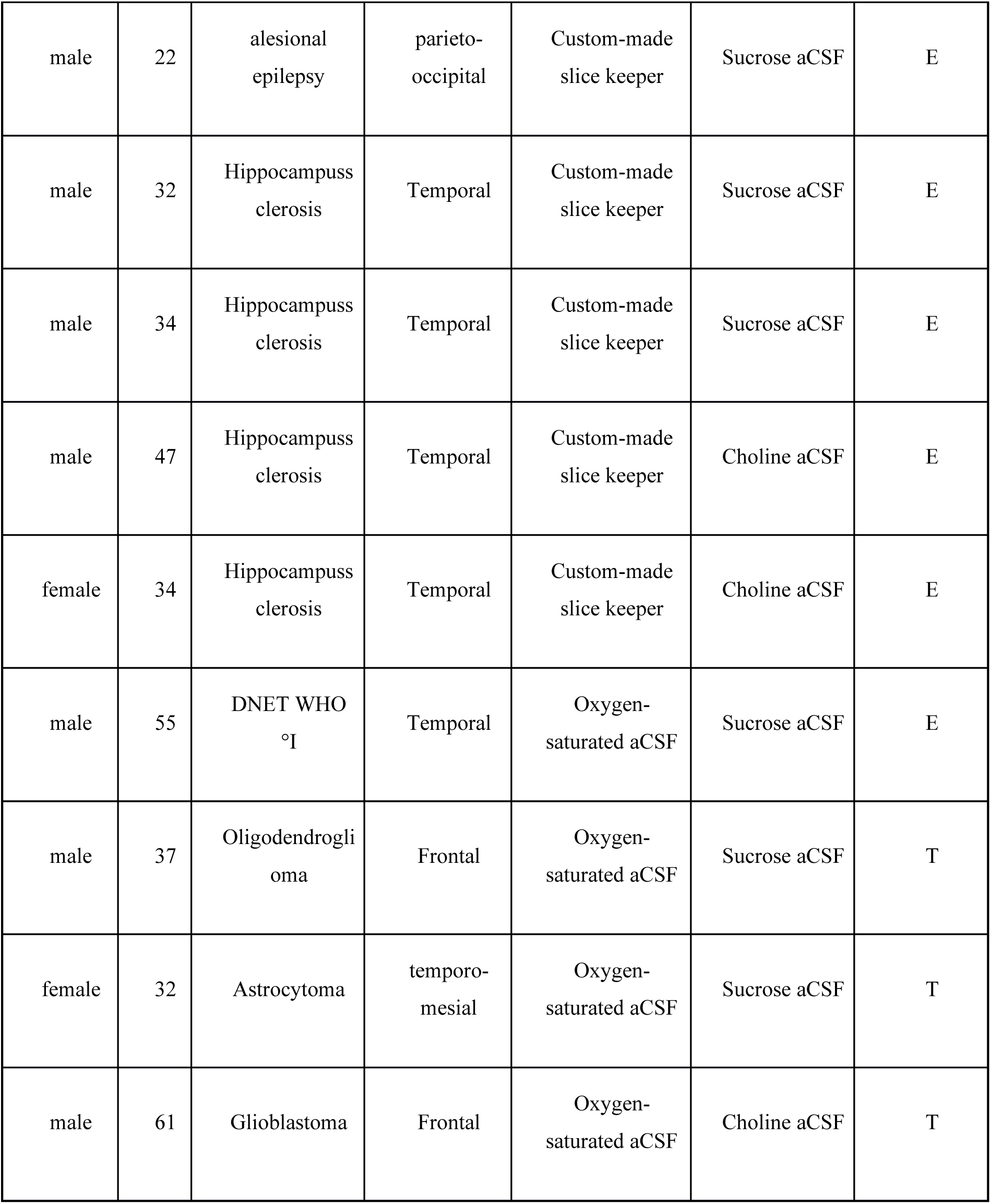
Patient details of brain surgery samples. In this study, tissue samples from n=32 surgeries were used. Patient details on gender, age, diagnosis, resection area, transportation method, preparation solution and pathology of the tissue samples are given.

Transport-Related Effects on Human Cortical Neurons
| **Gender** | **Age** | **Diagnosis** | **Resection area** | **Transportation method** | **Preparation solution** | **Epilepsy (E)**<br>**Tumor (T)** |
| --- | --- | --- | --- | --- | --- | --- |
| female | 26 | Heterotopia | Temporal | No transport | Sucrose aCSF | E |
| female | 35 | Cavernoma | Temporal | No transport | Sucrose aCSF | E |
| female | 41 | Metastasis | Frontal | No transport | Sucrose aCSF | T |
| female | 39 | Unspecific pathological finding | Parietal | No transport | Sucrose aCSF | E |
| female | 75 | Glioma | Frontal | No transport | Sucrose aCSF | T |
| male | 48 | Glioma | Frontal | No transport | Sucrose aCSF | T |
| female | 62 | Cavernoma | Temporal | No transport | Sucrose aCSF | E |
| male | 56 | Glioma | Frontal | No transport | Sucrose aCSF | T |
| female | 69 | Metastasis | Frontal | No transport | Sucrose aCSF | T |
| male | 17 | Glioma | Temporal | No transport | Sucrose aCSF | T |
| female | 67 | Glioma | Parietal | No transport | Sucrose aCSF | T |
| male | 9 | DNET | Temporal | No transport | Sucrose aCSF | E |
| female | 41 | Hippocampuss<br>clerosis | Temporal | Custom-made<br>slice keeper | Sucrose aCSF | E |
| female | 43 | Glioma | Temporal | Custom-made<br>slice keeper | Sucrose aCSF | T |
| male | 65 | Glioma | Temporal | Custom-made<br>slice keeper | Sucrose aCSF | T |
| male | 44 | Glioma | Frontal | Custom-made<br>slice keeper | Sucrose aCSF | T |
| male | 45 | Hippocampuss<br>clerosis | Temporal | Custom-made<br>slice keeper | Sucrose aCSF | E |
| female | 74 | Meningioma | Temporal | Custom-made<br>slice keeper | Sucrose aCSF | E |
| male | 63 | Metastasis | Frontal | Custom-made<br>slice keeper | Sucrose aCSF | T |
| female | 48 | Hippocampuss<br>clerosis | Temporal | Custom-made<br>slice keeper | Sucrose aCSF | E |
| male | 34 | DGONC | fronto-basal | Custom-made<br>slice keeper | Sucrose aCSF | E |
| female | 37 | Hippocampuss<br>clerosis | Temporal | Custom-made<br>slice keeper | Sucrose aCSF | E |
| male | 65 | Glioma | Temporal | Custom-made<br>slice keeper | Sucrose aCSF | T |

|  |  |  |  |  |  |  |
| --- | --- | --- | --- | --- | --- | --- |
| male | 22 | alesional<br>epilepsy | parieto-<br>occipital | Custom-made<br>slice keeper | Sucrose aCSF | E |
| male | 32 | Hippocampuss<br>clerosis | Temporal | Custom-made<br>slice keeper | Sucrose aCSF | E |
| male | 34 | Hippocampuss<br>clerosis | Temporal | Custom-made<br>slice keeper | Sucrose aCSF | E |
| male | 47 | Hippocampuss<br>clerosis | Temporal | Custom-made<br>slice keeper | Choline aCSF | E |
| female | 34 | Hippocampuss<br>clerosis | Temporal | Custom-made<br>slice keeper | Choline aCSF | E |
| male | 55 | DNET WHO<br>°I | Temporal | Oxygen-saturated aCSF | Sucrose aCSF | E |
| male | 37 | Oligodendrogli<br>oma | Frontal | Oxygen-saturated aCSF | Sucrose aCSF | T |
| female | 32 | Astrocytoma | temporo-<br>mesial | Oxygen-saturated aCSF | Sucrose aCSF | T |
| male | 61 | Glioblastoma | Frontal | Oxygen-saturated aCSF | Choline aCSF | T |

### Transportation of slices

To transport slices from RWTH Aachen University Hospital to Research Centre Juelich, a custom-made transportation system including a custom-designed slice keeper and a portable carbogen gas bottle (**Fig. 1B)** or a 50 ml tube with saturated carbogen was used. No significant difference in intrinsic properties of off-site recorded neurons between the two transportation methods were found (**Fig. 1-1**). The transportation time of the slices was between 30-40 minutes, covering approximately 38 kilometers. During transportation, slices were kept at a temperature of 20-25°C in aCSF and constantly oxygenated with 95% O_2_ and 5% CO_2_. Upon arrival, slices were transferred into a bubble chamber filled with carbogenated aCSF identical to the on-site set-up in Aachen **(Fig.1)**.

### Whole cell patch clamp recordings

Whole-cell recordings were performed in acute slices on-site (RWTH Aachen University Hospital) and off-site (Research Centre Juelich). During the patch-clamp recordings slices were continuously perfused (perfusion speed ∼ 5 ml/min) with aCSF, bubbled with carbogen gas, and maintained at 32–33°C. Patch pipettes were pulled from thick-walled borosilicate glass capillaries and filled with an internal solution containing (in mM): 135 K-gluconate, 4 KCl, 10 HEPES, 10 phosphocreatine, 4 Mg-ATP, and 0.3 GTP (pH 7.4 290–300 mOsm). Whole-cell patch-clamp recordings were performed using patch pipettes with a resistance of 5 – 10 MΩ and obtained using an EPC10 amplifier (HEKA). Signals were sampled at 10 kHz, filtered at 2.9 kHz using the Patchmaster software (HEKA), and later analyzed off-line using Igor Pro software (Wavemetrics). Biocytin was added to the internal solution at a concentration of 3–5 mg/ml for post-hoc histology. Neurons were visualized using either Dodt gradient contrast or infrared differential interference contrast microscopy. Putative pyramidal cells and interneurons were differentiated based on the appearance of cell bodies, and their AP firing pattern during the recordings and after post hoc histological staining by their morphological appearance.

### Histological processing

After recordings, brain slices containing biocytin-filled neurons were fixed for at least 24 h at 4 °C in 100 mM phosphate buffer solution (PBS, pH 7.4) containing 4% paraformaldehyde (PFA), following a published protocol (Marx et al., 2012). After rinsing several times in 100 mM PBS, slices were treated with 1% H_2_O_2_ in PBS for about 20 min to reduce any endogenous peroxidase activity. Slices were rinsed repeatedly with PBS and then incubated in 1% avidin-biotinylated horseradish peroxidase (Vector ABC staining kit, Vector Lab. Inc.) containing 0.1% Triton X-100 for 1 h at room temperature. The reaction was catalyzed using 0.5 mg/ml 3,3-diaminobenzidine (DAB; Sigma-Aldrich) as a chromogen. Subsequently, slices were rinsed with 100 mM PBS, followed by slow dehydration with ethanol in increasing concentrations, and finally in xylene for 2–4 h. Following that, slices were embedded using Eukitt medium (Otto Kindler GmbH).

### Morphological reconstructions

Using Neurolucida® software (MBF Bioscience, Williston, VT, USA), morphological reconstructions of biocytin-filled human L2/3 pyramidal cells and fast-spiking interneurons were made at a magnification of 1000-fold (100-fold oil-immersion objective and 10-fold eyepiece) on an upright microscope. Neurons were selected for reconstruction based on the quality of biocytin labeling when background staining was minimal. Neurons with major truncations due to slicing were excluded. Embedding using Eukitt medium reduced the fading of cytoarchitectonic features and enhanced contrast between layers. This allowed the reconstruction of different layer borders along with the neuronal reconstructions. Furthermore, the position of soma and layers was confirmed by superimposing the Dodt gradient contrast or differential interference contrast images taken during the recording. The tissue shrinkage was corrected using correction factors of 1.1 in the x–y direction and 2.1 in the z direction (Marx et al., 2012). In addition to reconstructing dendritic branches, spines located on dendrites of pyramidal cells were counted. For spine counting, approximately 100 µm-length branches belonging to basal, apical oblique and apical tuft dendrites were selected for each pyramidal cell. Analysis of 3D-reconstructed neurons was done with Neurolucida® Explorer software (MBF Bioscience, Williston, VT, USA).

### Data analysis

Custom-written macros for Igor Pro 6 (WaveMetrics) were used to analyze the recorded electrophysiological signals. The resting membrane potential (Vm) of the neuron was measured directly after the breakthrough to establish the whole-cell configuration with no current injection. The input resistance was calculated as the slope of the linear fit to the current-voltage relationship. For the analysis of single spike characteristics such as threshold, amplitude, and half-width, a step size increment of 10 pA for current injection was applied to ensure that the AP was elicited very close to its rheobase current. The spike threshold was defined as the point of start of acceleration of the membrane potential using the second derivative (d^2^V/dt^2^), that is, using 3x standard deviation of d^2^V/dt^2^ as the cut-off point. The spike amplitude was calculated as the difference in voltage from the AP threshold to the peak during depolarization. The spike half-width was determined as the time difference between the rising phase and the decaying phase of the spike at half-maximum amplitude.

The spontaneous synaptic activity was analyzed using the program SpAcAn (https://www.wavemetrics.com/project/SpAcAn). EPSPs and LRDs were distinguished by dramatic differences in event amplitude and decay time. A threshold of 0.2 mV was set manually for detecting EPSP events while a threshold of 3 mV was set for detecting LRDs.

### Statistical analysis

Data was either presented as box plots (n ≥ 10) or as bar histograms (n < 10). For box plots, the interquartile range (IQR) is shown as box, the range of values within 1.5*IQR is shown as whiskers and the median is represented by a horizontal line in the box; for bar histograms, the mean ± SD is given. Wilcoxon Mann-Whitney U-test was performed to assess the difference between individual clusters. A χ-square test was performed to assess differences between proportions. Statistical significance was set at p < 0.05, and n indicates the number of neurons analyzed.

## Results

### Recordings at the off-site laboratory

In the first set of experiments, we investigated the viability of cortical neurons in slices for electrophysiological recordings after being transported to the off-site laboratory as reported before (Mittermaier et al. 2024). We confirmed that human cortical neurons at the off-site laboratory remained viable for up to 60 hours and showed only minor changes in passive membrane properties, e.g. a decrease in input resistance but no significant change in their firing properties (**Fig. 2A**) or their post-hoc somatodendritic morphology (**Fig. 2B**) when compared between Day 1, Day2 or Day 3. However, we identified significant changes in sEPSP frequency and amplitude (**Fig. 2-1**). The sEPSP frequency significantly decreased in recordings obtained at Day 2 and Day 3 compared to Day 1, while the frequency was reduced at Day 2 compared to Day 1 and Day 3. Furthermore, a limited number of paired recordings were performed: In total, one L2/3 pyramidal-pyramidal pair, one L2/3 pyramidal-interneuron pair (**Fig. 3**), one L2/3 interneuron-pyramidal pair and one L2/3 pyramidal-interneuron reciprocal pair were recorded from transported human cortical brain slices. These paired recordings suggest that synaptically connected neurons can still be recorded after tissue transportation as previous reported (Mittermaier et al. 2024).

**Fig. 2.**
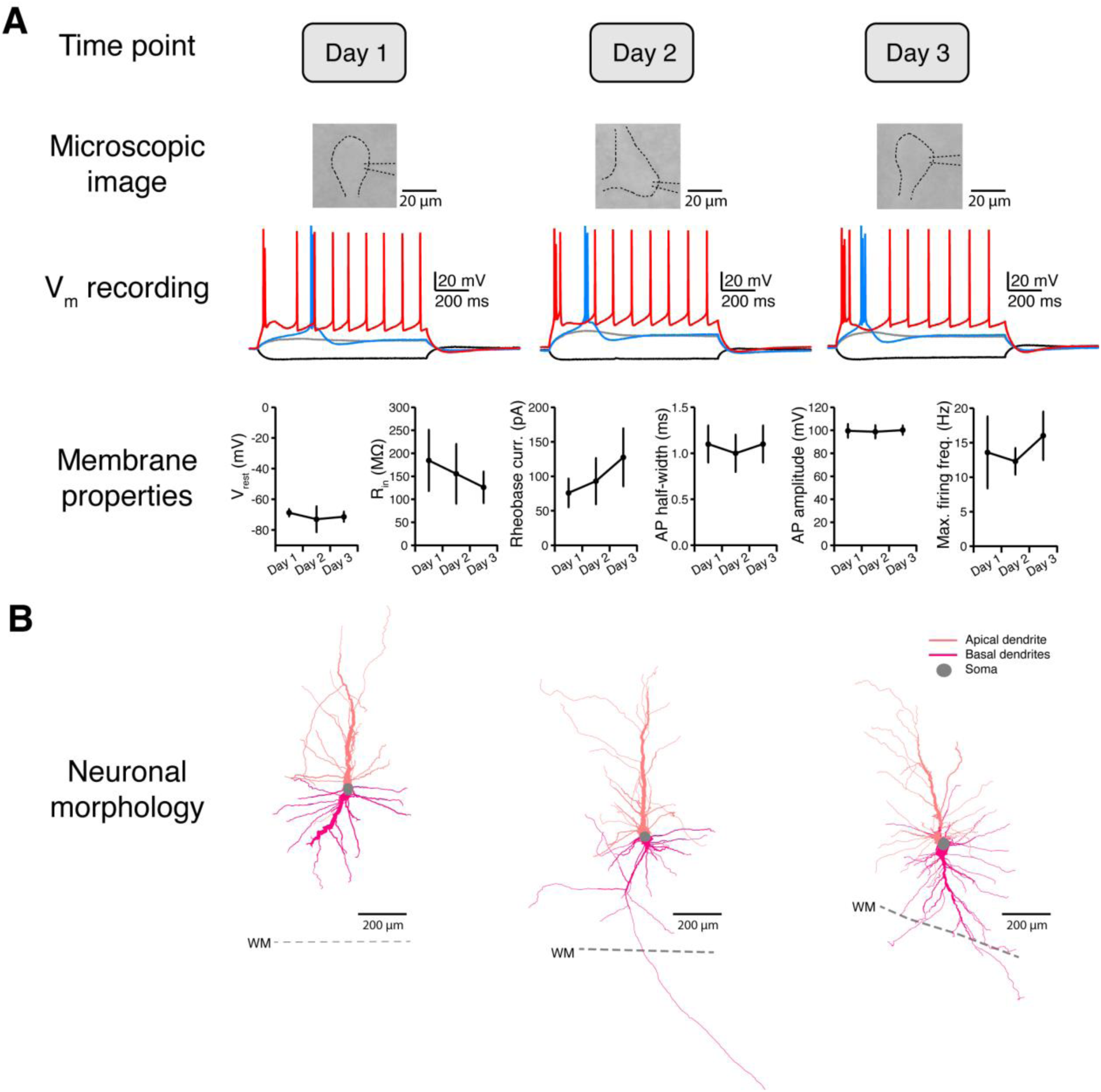
Prolonged (∼ 60 hrs) survival of human cortical neurons after transportation. (A) Representative patch-clamp recordings from cortical neurons in *ex vivo* slices on Day 1, Day 2, and Day 3 after slice preparation. Current-clamp recordings show the firing response to current injections. Bottom panel shows changes in intrinsic membrane properties over three days of recordings: n=5 for Day 1, n=7 for Day 2, and n=4 for Day 3. (B) Morphological reconstructions of three recorded human cortical neurons of which recordings are shown above. Cell bodies of these neurons were in the deep cortical layers near the white matter (WM).

**Fig. 3.**
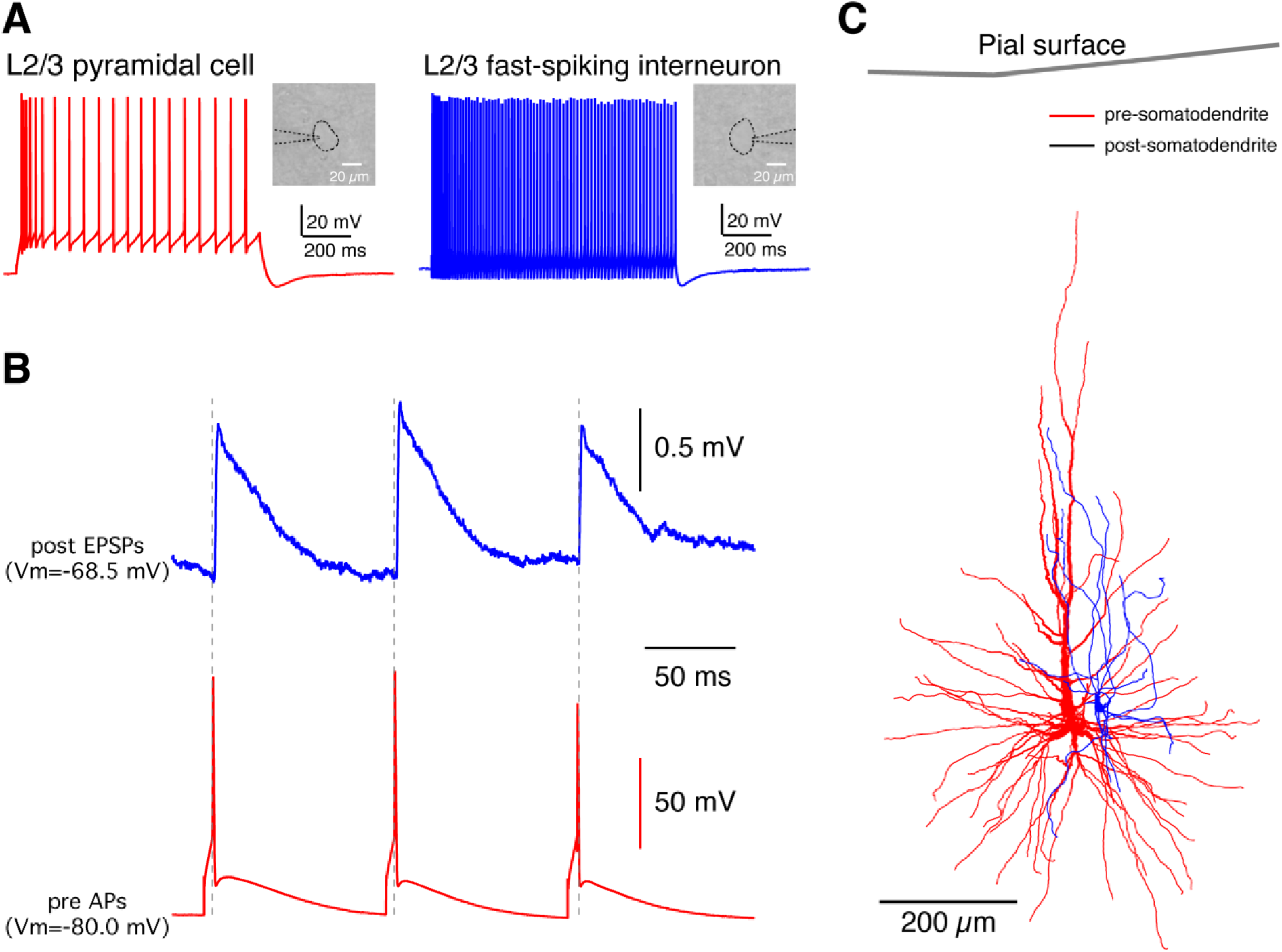
Paired recordings from synaptically coupled neurons in acute human cortical slices directly after transportation. (A) Firing patterns of pre– and postsynaptic neurons for synaptically coupled L2/3 pyramidal-FS interneuron pair. Inserts: DIC images of recorded neurons. (B) Postsynaptic EPSPs (top) were evoked by presynaptic APs (bottom).(C) Morphological reconstructions of the somatodendritic domain of pre– and postsynaptic neurons based on the post-hoc biocytin staining.

### Off-site recordings show different intrinsic properties after transportation

To determine whether acute human brain slices can be reliably used for electrophysiological patch-clamp recordings in a laboratory located off-site from the clinic where the slices were prepared, and to assess if transport affects the properties of recorded neurons, we performed either on-site or off-site recordings or parallel on– and off-site recordings (n=32 surgeries, see Table 1). Slices were recorded either at the RWTH Aachen University Hospital (on-site) or at the Research Center Juelich (off-site).

In total, this study comprised 200 neurons, with 133 neurons recorded on-site and 67 neurons recorded off-site. First, we compared the intrinsic firing properties of L2/3 pyramidal cells and fast-spiking (FS) interneurons recorded in parallel on-site and off-site. Only neurons with a stable resting membrane potential below –60 mV and a series resistance smaller than 40 MΩ were included in the analysis to ensure a high recording quality.

L2/3 pyramidal cells (n = 10 on-site and n = 11 off-site) at both recording sites exhibited the characteristic firing pattern of regular action potential firing with spike frequency adaptation (**Fig. 4A**), as previously described (Avoli 1994). However, several intrinsic properties of these neurons showed significant differences between on-site and off-site recordings (**Fig. 4B**). Specifically, pyramidal cells recorded at the off-site laboratory were slightly more depolarized, exhibiting a less negative resting membrane potential compared to those recorded on-site. Additionally, off-site pyramidal cells had a significantly lower rheobase current and a higher maximal firing frequency compared to their on-site counterparts. In conclusion, neurons recorded off-site were more excitable and required less current input to reach the threshold for action potential generation.

**Fig. 4.**
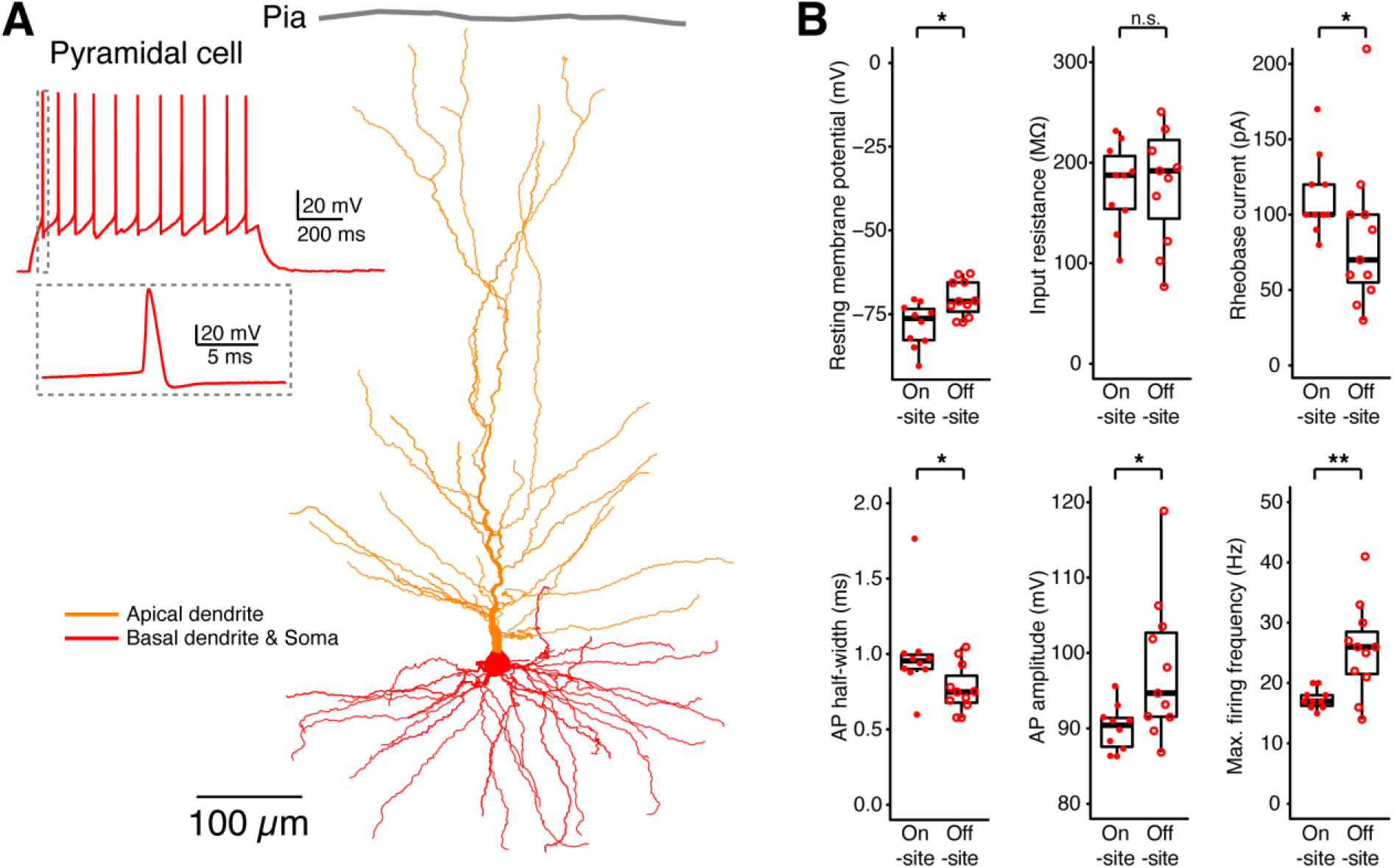
Comparison of the electrophysiological properties between on– and off-site recorded human cortical L2/3 pyramidal cells. (A) Typical firing pattern (regular firing) and the morphological reconstruction of a human cortical L2/3 pyramidal cell. (B) Comparison of electrophysiological properties of pyramidal cells recorded on-site (n=10) and off-site (n=11). n.s. p≥0.05, * p<0.05, ** p<0.01 for the Wilcoxon-Mann-Whitney two-sample rank test.

These results suggest that while the overall firing capability of L2/3 pyramidal cells is independent of recording location, specific intrinsic properties, such as resting membrane potential and rheobase current, are influenced by the transportation process. Similarly, FS interneurons (n = 11 on-site and n = 10 off-site) recorded at both locations demonstrated the expected characteristic high-frequency firing with narrow action potentials and no spike frequency adaptation (**Fig. 5A**). However, significant differences were observed in specific action potential parameters between the on-site and off-site recordings (**Fig. 5B**).

**Fig. 5.**
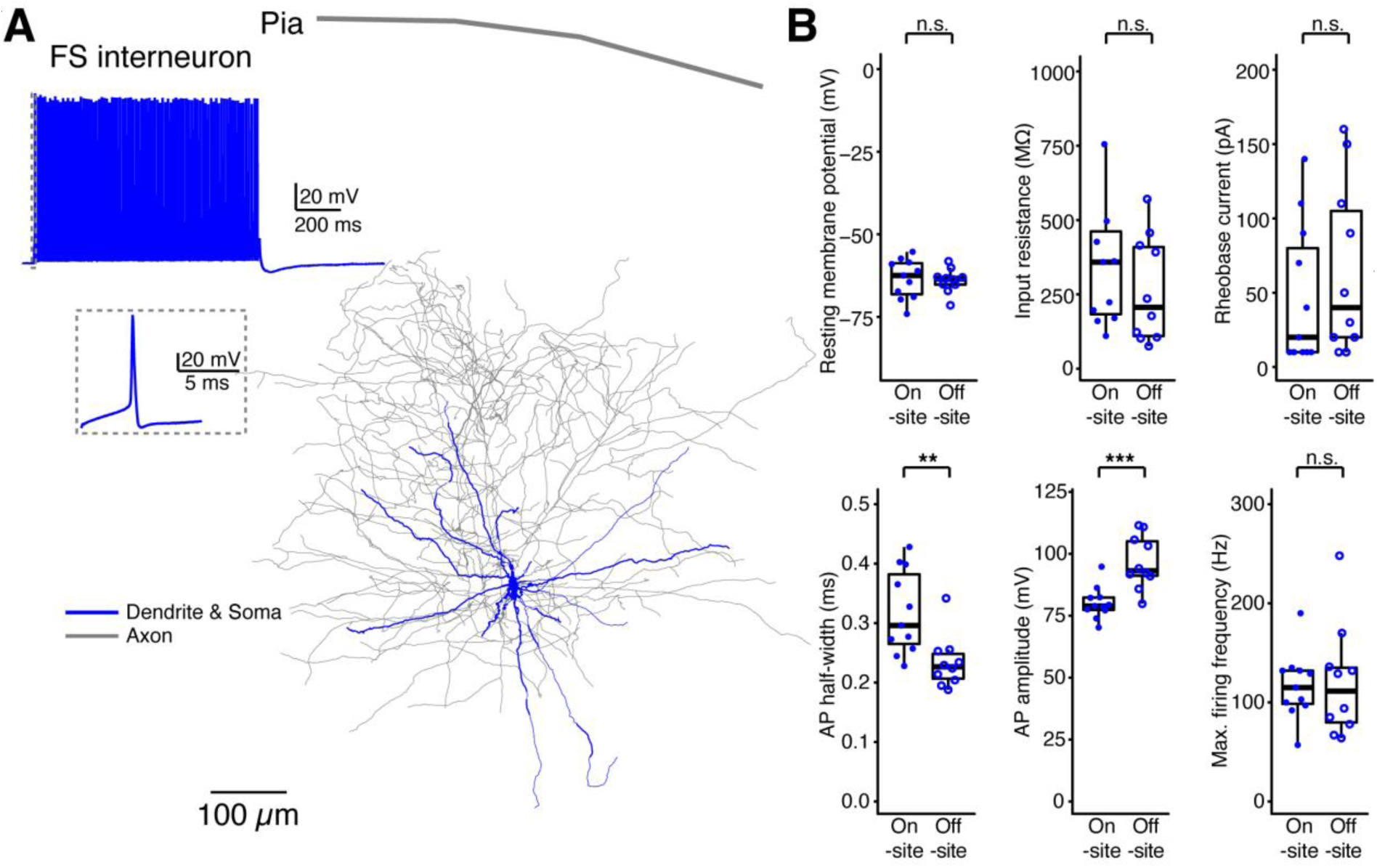
Comparison of electrophysiological properties of human fast-spiking (FS) interneurons recorded on– and off-site. (A) Representative firing pattern and morphological reconstruction of a human cortical L2/3 FS interneuron. (B) Comparison of electrophysiological properties of FS interneurons recorded on-site (n=11) and off-site (n=10). n.s. p≥0.05, ** p<0.01, *** p<0.001 for the Wilcoxon Mann-Whitney U-test.

### Comparison of spontaneous EPSP frequency and amplitude between on-site and off-site recorded human cortical neurons

To investigate whether the transport of human cortical slices to an off-site laboratory affects synaptic properties, we compared the frequency and amplitude of spontaneous excitatory postsynaptic potentials (sEPSPs) recorded in L2/3 pyramidal cells and interneurons in parallel on-site and off-site. Only cells from surgeries were recordings were performed both on– and off-site were included in this analysis. L2/3 pyramidal cells recorded on-site (Fig. 6A upper trace) and off-site (**Fig. 6A** lower trace) exhibited sEPSP activity (**Fig. 6A**, left).

**Fig. 6.**
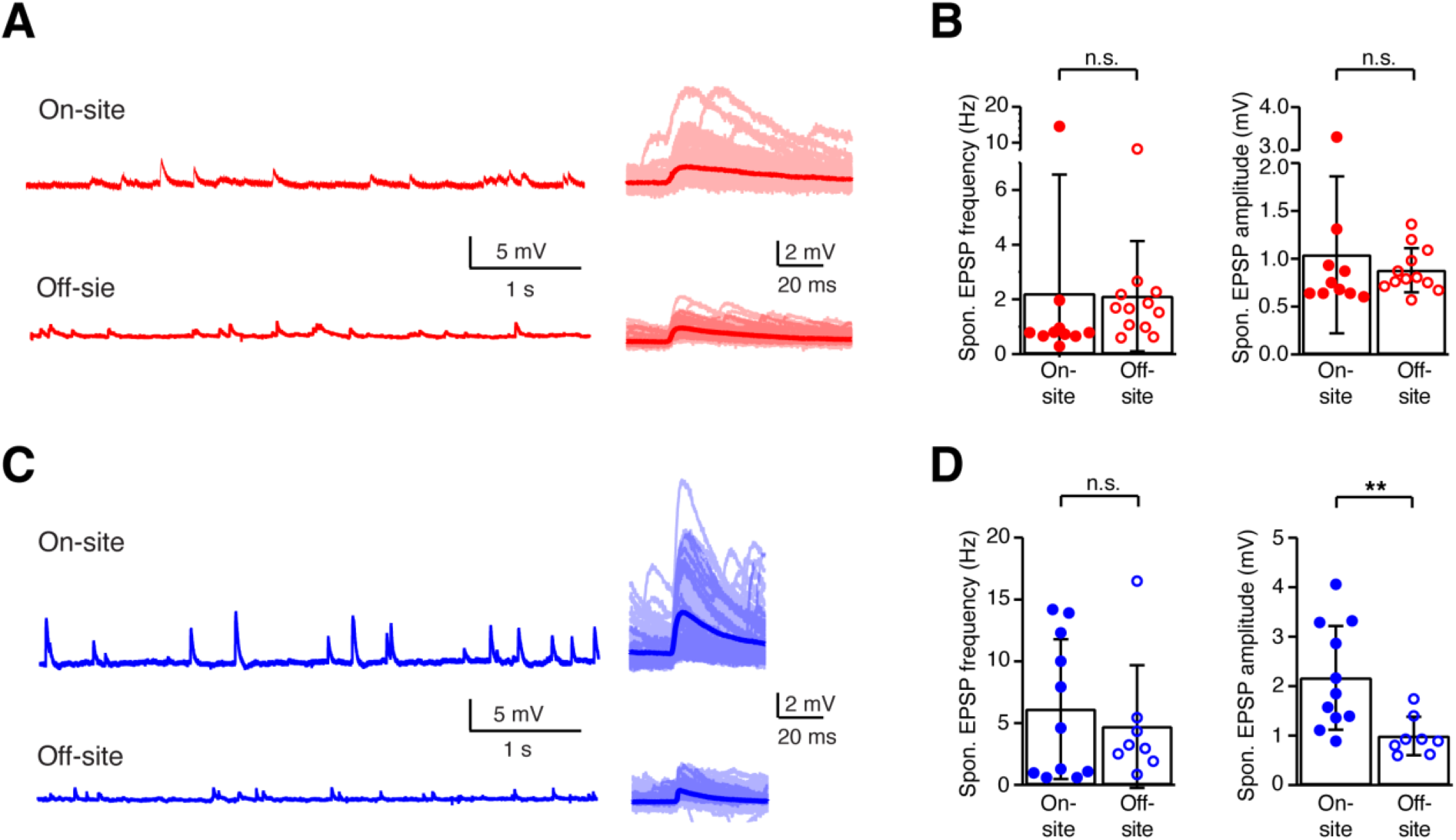
Comparison of spontaneous EPSP frequency and amplitude between on-site and off-site recorded human cortical neurons. (A) Left: Representative Vm traces of human cortical L2/3 pyramidal cells recorded on-site (upper trace) and off-site (lower trace). Right: Overlay of single spontaneous EPSPs (sEPSPs) and their average for visual comparison. (B) Comparison of sEPSP frequency (left) and amplitude (right) in pyramidal cells recorded on-site (n = 10) versus off-site (n = 12). No significant differences were observed in either EPSP frequency or amplitude between the two recording locations (n.s., p ≥ 0.05, Wilcoxon-Mann-Whitney two-sample rank test). (C, D) Similar analysis as in (A, B) but for human cortical L2/3 interneurons recorded on-site (n = 11) and off-site (n = 8). While there were no significant differences in sEPSP frequency between on-site and off-site recordings, a significant decrease in sEPSP amplitudes was observed in off-site recorded interneurons in comparison to those recorded on-site (p < 0.01, Wilcoxon-Mann-Whitney two-sample rank test.

Quantitative analysis confirmed that there were no significant differences in the frequency or amplitude of sEPSPs between pyramidal cells recorded on-site (n = 10) and those recorded off-site (n = 12) (**Fig. 6B**). Similarly, the analysis of L2/3 interneurons recorded on-site (n = 11) and off-site (n = 8) revealed no significant differences in sEPSP frequency between the two groups (p ≥ 0.05) (**Fig. 6C** and **6D**, left). However, a significant decrease in sEPSP amplitude was observed in interneurons recorded off-site compared to those recorded on-site (p < 0.01, Wilcoxon Mann-Whitney U-test) (**Fig. 6D**, right). This reduction in sEPSP amplitude suggests that the synaptic efficacy of excitatory connections to interneurons is particularly susceptible to the effects of transport. Overall, these findings indicate that while the frequency of sEPSPs remains consistent between on-site and off-site recordings for both pyramidal cells and interneurons, the amplitude of EPSPs in interneurons was significantly reduced when recordings are conducted off-site compared to off-site recordings.

This suggests that transport conditions may affect synaptic transmission, particularly in excitatory-to-inhibitory circuits, potentially impacting their ability to modulate network activity effectively.

### Comparison of LRD occurrence in brain slice samples recorded on-site and off-site

Next, we recorded the spontaneous activity of neurons in current-clamp mode to determine the presence of spontaneous large rhythmic depolarizations (LRDs), which have been described recently (Yang et al., 2024). At the on-site laboratory, 10% of the recorded neurons exhibited spontaneous LRDs, whereas only 3% of the neurons recorded at the off-site laboratory displayed spontaneous LRDs (**Fig. 7A-C**). Interestingly, neurons displaying spontaneous LRDs that were recorded at the off-site laboratory were located significantly deeper in the slice than those neurons showing no LRD: the depth from soma to the slice surface is 92.5 ± 10.6 µm (n = 2) for LRD+ neurons and 74.7 ± 16.0 µm (n = 30) for LRD− neurons. Additionally, in a subset of neurons, we used norepinephrine (NE, 10 or 30 µM) as a pharmacological stimulus to elicit LRDs (Yang et al., 2024). We found that 21% of the neurons recorded at the on-site laboratory responded with LRDs following NE application, while only 7% of the neurons recorded off-site exhibited a similar response (**Fig. 7D-E**). Hence, the incidence of LRDs was significantly reduced at the off-site laboratory, both in the absence and presence of NE induced modulation. This indicates that synaptic function and network activity are vulnerable to transport-related effects.

**Fig. 7.**
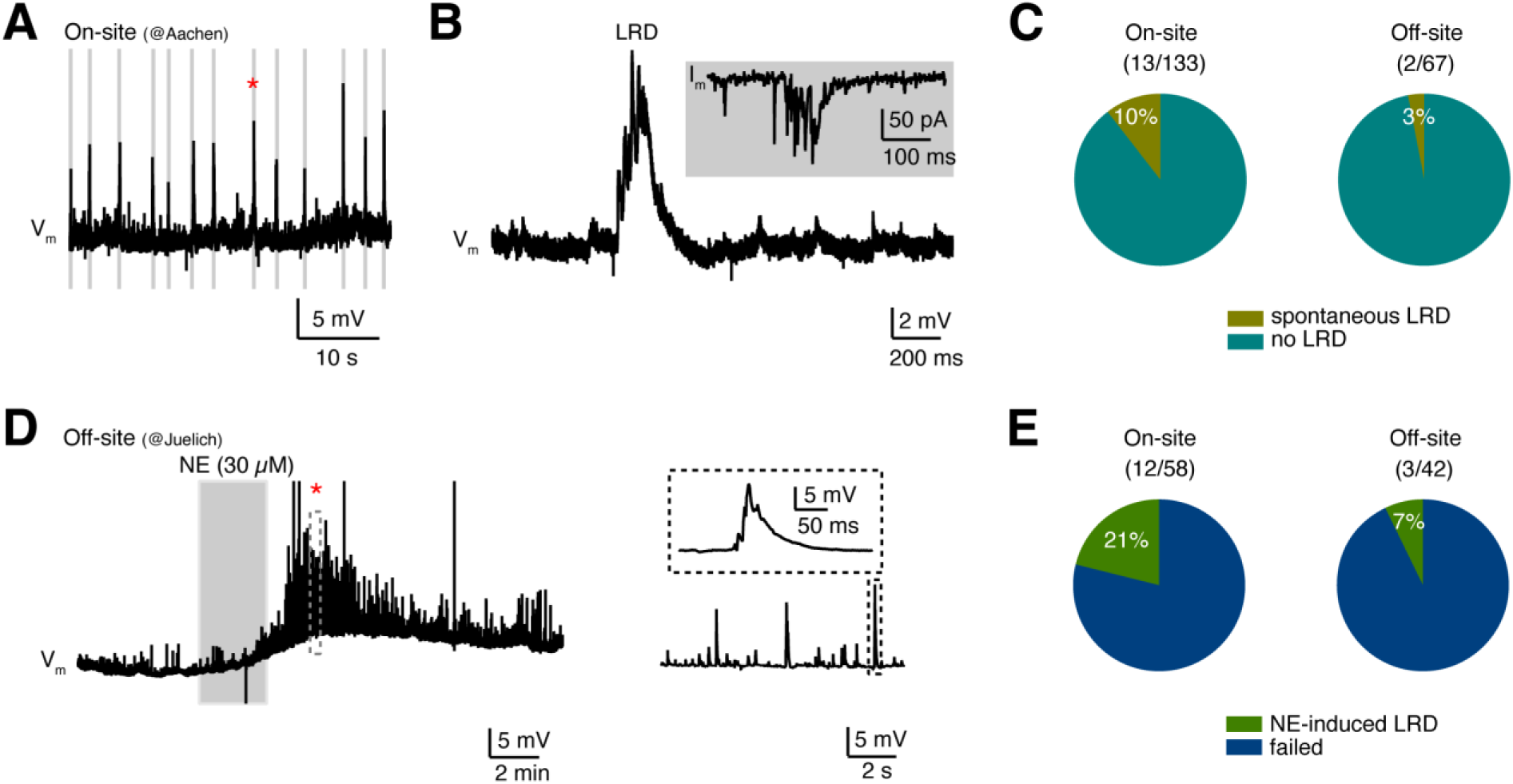
Self-organized network activity was predominantly observed in human cortical neurons recorded on-site. (A) Vm recordings from a human cortical L2/3 interneuron demonstrating spontaneously occurring large rhythmic depolarizations (LRDs). This neuron was recorded on-site at the Aachen laboratory. (B) A single LRD marked by an asterisk in (A) is enlarged, which is composed of several EPSPs at the rising phase. Inset shows: Voltage-clamp recording of a single LRD from the same neuron. (C) Percentage of on– and off-site recorded neurons displaying spontaneous LRDs. P = p = 0.043 for χ-square test without Yates correction and therefore significant (n.s. p≥0.05. (D) LRDs were induced by bath application of noradrenaline (NE, 10 or 30 μM) in neurons that showed no spontaneous LRDs. (E) Percentage of neurons exhibiting NE-induced LRDs recorded on– and off-site. P = 0.031 for χ-square test without Yates correction and therefore significant (n.s., p≥0.05).

### Post-hoc morphology of pyramidal cells and interneurons

The post-hoc morphological analysis of pyramidal cells and interneurons recorded either on-site or off-site revealed no significant differences between the two groups in overall dendritic structure, including dendritic length and branching pattern (**Fig. 8A and 8B**, and **Table 3**). This indicates that despite the observed differences in electrophysiological properties, the transport of brain slices to an off-site laboratory does not affect the overall structural integrity of the neurons.

**Fig. 8.**
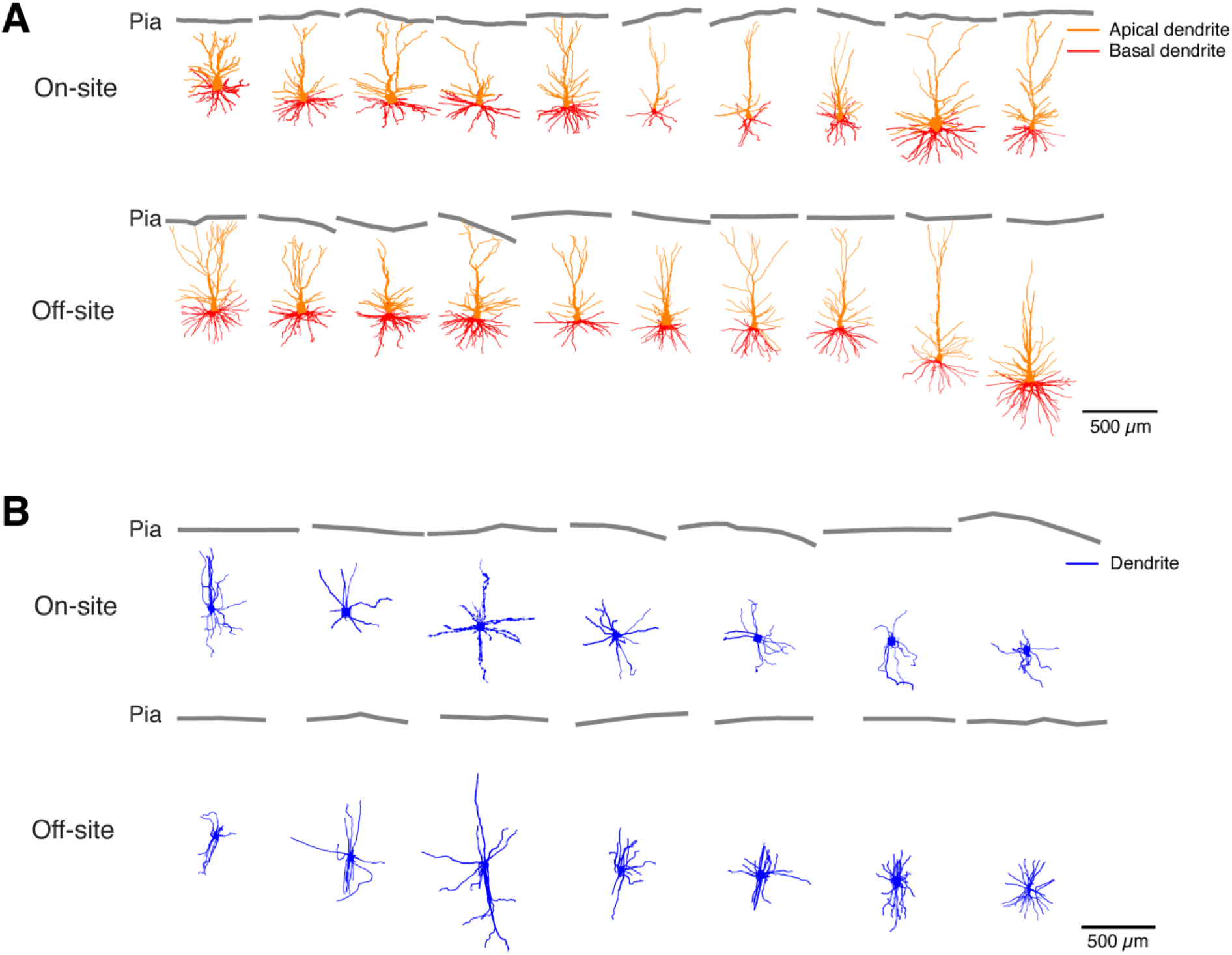
Comparison of dendritic morphologies between on– and off-site recorded human cortical neurons. (A) Morphological reconstructions of human cortical L2/3 pyramidal cells recorded on-site (top, n=10) and off-site (bottom, n=10). (B) Morphological reconstructions of human cortical L2/3 FS interneurons recorded on-site (top, n=7) and off-site (bottom, n=7). No significant difference in somatodendritic morphologies was found between on– and off-site recorded neurons.

**Table 2.** Comparison of electrophysiological properties between On– and Off-site recorded human cortical neurons. The P-value was calculated using the non-parametric Wilcoxon-Mann-Whitney two-sample rank test. Data was presented as mean ± standard deviation.

|  | L2/3 pyramidal cells |  |  | L2/3 FS interneurons |  |  |
| --- | --- | --- | --- | --- | --- | --- |
|  | On-site<br>(n=10) | Off-site<br>(n=11) | P value | On-site<br>(n=11) | Off-site<br>(n=10) | P value |
| *Passive* |  |  |  |  |  |  |
| **Vrest (mV)** | -78.2 ± 6.6 | -70.4 ± 5.4 | 0.024 | -63.5 ± 5.9 | -64.0 ± 3.7 | 0.70 |
| **Rin (MΩ)** | 177.5 ± 41.7 | 192.4 ± 82.7 | 0.70 | 432.7 ± 400.8 | 265.4 ± 178.0 | 0.31 |
| **Tau (ms)** | 17.4 ± 5.0 | 22.3 ± 5.5 | 0.085 | 13.7 ± 4.8 | 10.3 ± 2.6 | 0.15 |
| **Sag (%)** | 8.8 ± 10.8 | 14.5 ± 13.6 | 0.31 | 24.9 ± 16.2 | 18.4 ± 7.6 | 0.43 |
| *Single AP* |  |  |  |  |  |  |
| **Rheobase current (pA)** | 112.0 ± 26.6 | 84.5 ± 50.1 | 0.042 | 47.3 ± 47.6 | 65.0 ± 58.2 | 0.34 |
| **AP threshold (mV)** | -41.3 ± 4.9 | -41.2 ± 3.7 | 0.97 | -41.2 ± 4.4 | -40.7 ± 4.1 | 1.03 |
| **AP half-width (ms)** | 1.0 ± 0.3 | 0.8 ± 0.2 | 0.043 | 0.32 ± 0.07 | 0.23 ± 0.04 | 2.1×10<sup>-3</sup> |
| **AP amplitude (mV)** | 90.0 ± 3.0 | 97.8 ± 9.3 | 0.016 | 80.1 ± 6.5 | 96.6 ± 10.7 | 7.9×10<sup>-4</sup> |
| **AHP amplitude (mV)** | 16.1 ± 4.8 | 14.2 ± 2.6 | 0.35 | 25.4 ± 5.0 | 27.4 ± 2.9 | 0.35 |
| *Repetitive firing* |  |  |  |  |  |  |
| **Max. firing frequency (Hz)** | 17.4 ± 1.6 | 25.5 ± 7.6 | 7.2×10<sup>-3</sup> | 116.5 ± 33.7 | 120.3 ± 56.7 | 0.82 |
| **Slope of F-I curve (APs/100 pA)** | 16.7 ± 5.0 | 14.0 ± 4.8 | 0.25 | 56.1 ± 35.8 | 49.0 ± 16.9 | 0.97 |

**Table 3:** Comparison of morphological properties between On– and Off-site recorded human cortical neurons. The p-value was calculated using the non-parametric Wilcoxon-Mann-Whitney two-sample rank test. Data is presented as mean ± standard deviation. N.A., not applicable.

|  | **L2/3 pyramidal cells** |  |  | **L2/3 FS interneurons** |  |  |
| --- | --- | --- | --- | --- | --- | --- |
|  | **On-site**<br>**(n=10)** | **Off-site**<br>**(n=10)** | **P value** | **On-site**<br>**(n=7)** | **Off-site**<br>**(n=7)** | **P value** |
| **No. of basal dendrites** | 5.7 ± 1.7 | 4.9 ± 0.7 | 0.34 | 6.3 ± 1.5 | 6.3 ± 1.2 | 0.89 |
| **No. of basal dendrite branches** | 27.8 ± 9.5 | 31.3 ± 9.1 | 0.50 | 16.1 ± 4.7 | 18.3 ± 4.3 | 0.51 |
| **Total length of basal dendrites (mm)** | 4.9 ± 1.8 | 5.2 ± 1.6 | 0.92 | 4.3 ± 1.3 | 4.6 ± 1.3 | 0.63 |
| **Horizontal field span of basal dendrites (mm)** | 45.3 ± 13.7 | 46.7 ± 7.1 | 0.84 | 49.0 ± 14.6 | 46.8 ± 19.7 | 0.84 |
| **Vertical field span of basal dendrites (μm)** | 29.1 ± 6.0 | 34.8 ± 10.5 | 0.29 | 59.8 ± 16.4 | 68.6 ± 30.0 | 0.73 |
| **No. of apical dendrite branches** | 22.6 ± 7.8 | 24.9 ± 9.1 | 0.89 | N.A. | N.A. | N.A. |
| **Total length of apical dendrite (mm)** | 5.1 ± 1.7 | 5.2 ± 1.8 | 0.92 | N.A. | N.A. | N.A. |
| **Horizontal field span of apical dendrite (mm)** | 41.5 ± 11.5 | 41.2 ± 4.2 | 0.84 | N.A. | N.A. | N.A. |

However, a detailed analysis of dendritic spine densities for L2/3 pyramidal cells (**Fig. 9A**), which are critical for synaptic function and plasticity, revealed significant differences between the two recording sites. Specifically, we found that the spine densities in the apical oblique and apical tuft dendrites were significantly reduced in pyramidal cells recorded off-site compared to those recorded on-site (**Fig. 9C**).

**Fig. 9.**
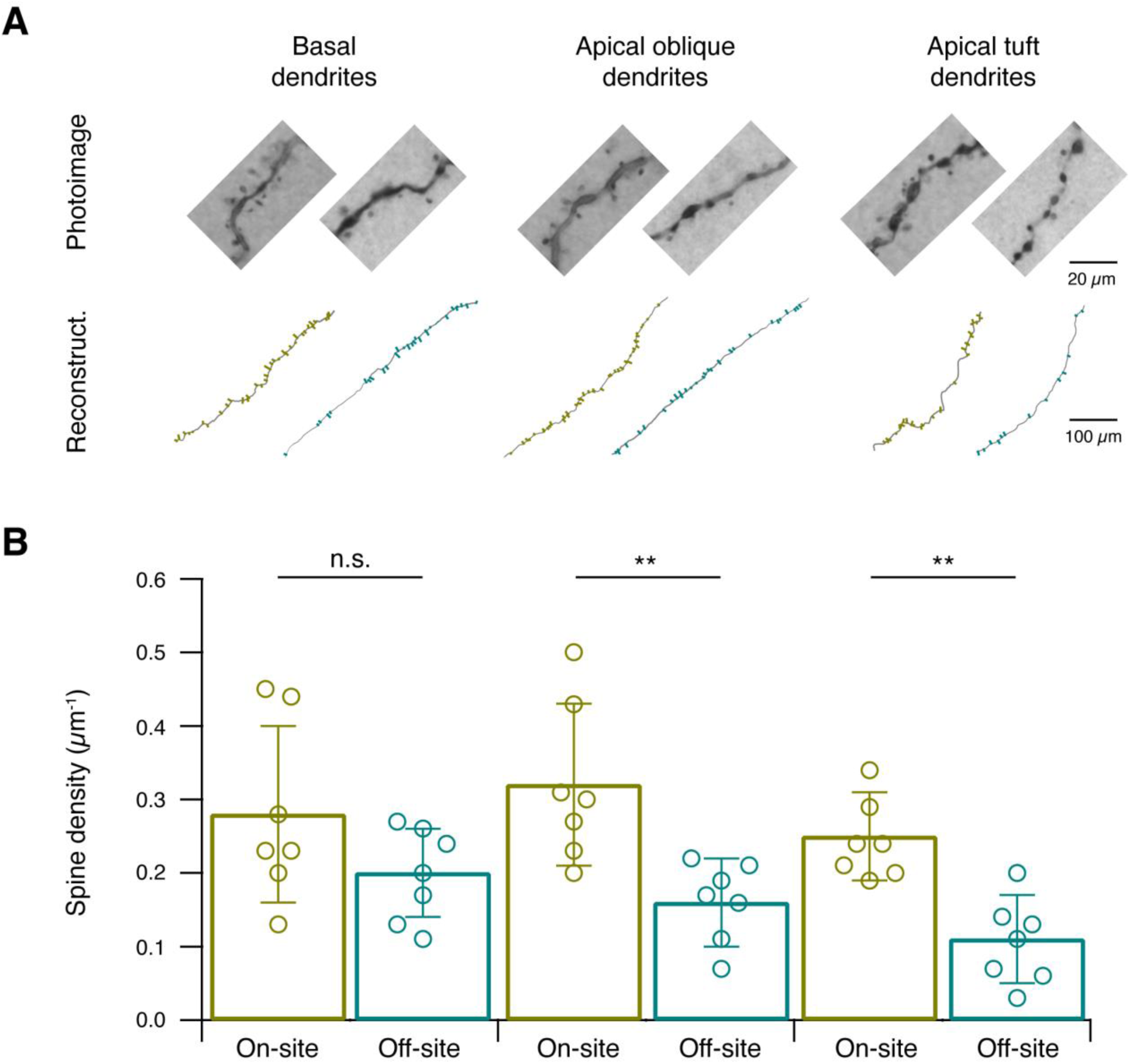
Comparison of spine densities between on– and off-site recorded human cortical L2/3 pyramidal cells. (A) Representative images and reconstructions of spines at basal, apical oblique, and apical tuft dendrites for on– and off-site recorded L2/3 pyramidal cells after biocytin staining. (B) Comparison of spine densities at the three dendritic compartments left: basal dendrites, middle: Apical oblique and right Apical tuft for on-site (n=7) and off-site (n=7) recorded L2/3 pyramidal cells. Significant reductions in spine densities were found at the apical oblique and apical tuft dendrites for off-site recorded L2/3 pyramidal cells.

In contrast, no significant differences were observed in the basal dendritic compartments of the two groups. These findings suggest that while the overall dendritic and axonal architecture remains intact, the finer synaptic structures, such as dendritic spines, are more susceptible to transport-related effects, potentially contributing to the observed alterations in synaptic efficacy and network activity.

## Discussion

This study provides a comprehensive analysis of the effects of transportation on the intrinsic and synaptic properties of human cortical neurons by comparing on-site and off-site recordings. While a successful transportation of human brain tissue for live cell recordings has been previously documented (Köhling et al., 1996; Mittermeier et al., 2024; Andersson, 2016), detailed investigations into subtle changes in morphological, electrophysiological, and synaptic network properties are required to assess whether transport affects tissue integrity and neuronal function. The results of our study indicate that neurons in brain slices transported for 30-40 minutes from the surgical resection site to an off-site laboratory already exhibit alterations in intrinsic, synaptic, and morphological properties.

### Alterations in intrinsic properties of neurons

Our data show that while the overall firing capabilities of L2/3 pyramidal cells and FS interneurons are preserved between on-site and off-site recordings, there are subtle yet significant differences in intrinsic properties. Off-site recorded pyramidal cells were slightly more depolarized and exhibited a lower rheobase, suggesting increased excitability and sensitivity to depolarizing inputs. These changes may be linked to mechanical stress during transportation, which prompts microglia to release reactive oxygen species and cytokines (Banati, 1993). Microglia and astrocytes both have been shown to release pro-inflammatory cytokines in response to insults and neuronal inflammation which was demonstrated to enhance neuronal excitability by increasing sodium channel currents (Vezzani 2015, Wang, 2007; Takashi, 2007). For FS interneurons, off-site recordings showed a narrower action potential (AP) half-width and an increased amplitude compared to on-site recordings. This might result from the washout of residual neuromodulators in the intercellular space during transport, as neuromodulators like norepinephrine, acetylcholine and dopamine can modulate AP amplitude and width (Qi & Feldmeyer, 2022, Kawaguchi and Shindou, 1998; Gorelova et al., 2002). These alterations could influence overall network dynamics and cortical information processing.

### Impact of transport on synaptic function and network events

The reduced network activity after transportation could have important implications for the study of human brain tissue, where intact network dynamics are crucial for understanding pathological and physiological underlying mechanisms (de la Prida 2019, Nimmrich 2015). Our analysis of spontaneous excitatory postsynaptic potentials (EPSPs) indicates that synaptic function is particularly vulnerable to transport-related effects. While the frequency of spontaneous EPSPs did not differ significantly between on-site and off-site recordings for both pyramidal cells and interneurons, there was a notable reduction in EPSP amplitudes in off-site recorded interneurons. This suggests that synaptic efficacy, particularly in excitatory-to-inhibitory circuits, may be compromised during transport. The decreased EPSP amplitude could reflect reduced presynaptic release probability or postsynaptic receptor sensitivity, especially as EPSPs in human pyramidal neurons have been shown to be largely NMDA receptor-dependent (Hunt 2023, Eyal 2018).

Furthermore, one of the most noteworthy findings of our study is the significant reduction in the occurrence of spontaneous LRDs and NE-induced LRDs in neurons recorded off-site in comparison to those recorded on-site. Specifically, only 3% of neurons recorded off-site exhibited spontaneous LRDs, compared to 10% on-site. Similarly, the percentage of neurons showing NE-induced LRDs was significantly lower off-site than on-site (21%). As LRDs are predominantly observed in human interneurons rather than in pyramidal cells and depend on the glutamatergic transmission (Yang et al. 2024), the lower EPSP amplitude found in off-site recorded interneurons may serve as a cautionary indicator of reduced LRD occurrence. These findings support the notion that transport to an off-site laboratory has an adverse impact on network-level events, possibly due to subtle changes in the synaptic and neuromodulatory milieu of neurons. It may be hypothesized that vibrations occurring during transport could therefore diminish the concentrations of neuromodulators, such as NE, in the intercellular space. This could, in turn, lead to a substantial influence on the occurrence of network events such as LRDs (Yang et al. 2024).

### Mechanisms underlying transport-induced changes

Transport-induced mechanical stress can have various detrimental effects on human brain tissue, largely due to its impact on the mechanical properties of the tissue. During transport, the brain slices are continuously exposed to vibrations, acceleration, deceleration, and sudden jolts, which can exert mechanical stress on the delicate structures. These forces can alter the mechanical properties of the brain tissue, which are determined not only by cellular components but also by the extracellular matrix (ECM). The ECM, comprising elements like collagen (Shulyakov et al., 2011; Budday et al., 2020), proteoglycans (Lotz and Loeser, 2012), and lipids (Mihai and Goriely, 2017), provides crucial structural and biochemical support, influencing tissue stiffness, elasticity, and resilience. The mechanical stress encountered during transport can alter these ECM components, leading to changes in tissue hydration, viscoelastic properties, and membrane fluidity, ultimately affecting cellular behaviors such as signaling pathways.

Increased mechanical stress during transport has already been shown to significantly alter neuronal properties, including reduced ATP content (Ahmed, 2000) and increased plasma membrane permeability (Geddes, 2003). Even mild mechanical stress can dramatically elevate the production of reactive oxygen and nitrogen species (ROS/RNS) (Arundine, 2004). Additionally, mechanical stress can lead to a reduction in dendritic spines due to NMDA receptor-mediated increases in postsynaptic calcium (Chen, 2015). This surge in calcium can additionally be triggered by ROS/RNS and could induce abnormal homeostatic responses, further altering neuronal properties (Higgins, 2010).

It is worth noting that changes in homeostatic plasticity can lead to synaptic scaling, which often results in a reduction in dendritic spine density (Moulin et al., 2019, 2022). The decrease in spine density, and consequently synapse number, particularly near the apical dendrites, may contribute further to disrupted network activity. Oxidative stress can also disrupt the cytoskeleton, especially actin (Allani, 2004), which plays a critical role in the formation and regulation of dendritic spines (Lei, 2016).

### Implications for experimental design, future research, and conclusion

The findings of this study highlight how transport-induced stress can lead to significant alterations in synaptic efficacy and network dynamics, notably the reduction in EPSP amplitudes and LRDs. These changes, which may stem from mechanical stress and its impact on the extracellular matrix and neuromodulator availability, can compromise the reliability of experimental data, especially in studies focused on synaptic plasticity or network events. On-site recordings, where transport-induced effects are minimized, are likely to yield more accurate and physiologically relevant results.

To mitigate these effects, future research should aim to refine transport protocols. Enhancing transport conditions—such as adjusting temperature, oxygenation, and medium composition— could preserve tissue integrity. For example, transporting tissue blocks instead of slices may reduce mechanical stress on individual cells and help retain neuromodulators. However, hypoxia during transport remains a challenge that may be alleviated through hypothermia, which has been shown to preserve tissue better (Antonic, 2014).

Moreover, the molecular mechanisms underlying transport-induced alterations, such as the disruption of ion channel function, synaptic vesicle release, receptor dynamics, and cytoskeletal integrity, need further investigation. Mechanical stress-induced production of reactive oxygen and nitrogen species (ROS/RNS), as well as disruptions in calcium signaling and the ECM, may play critical roles in these changes. By studying these pathways in greater detail, researchers can develop standardized protocols for tissue transport, enhancing the reproducibility and reliability of human brain studies across laboratories

In conclusion, while the firing capacity of human cortical neurons remains largely intact after transport, significant changes in neuronal excitability, synaptic efficacy, and network-level events underscore the importance of considering transport effects in experimental designs. By optimizing transportation protocols and exploring the molecular changes induced by mechanical stress, researchers can better preserve tissue integrity. This will ultimately improve the quality and reliability of human brain tissue research, advancing our understanding of cortical microcircuits and their role in both healthy and diseased states.

## CRediT authorship contribution statement

**Qi Guanxiao:** Conceptualization, Investigation, Formal Analysis, Data Curation, Visualization, Validation, Methodology, Writing – review & editing.

**Yang Danqing:** Investigation, Formal Analysis, Data Curation, Visualization, Validation, Methodology, Writing – review & editing, Conceptualization.

**Bak Aniella Vanessa:** Writing – original draft, Writing – review & editing, Validation, Methodology. Hucko Werner: Investigation, Methodology and Validation.

**Hussam Hamou:** Writing – review & editing, Validation.

**Delev Daniel:** Writing – review & editing, Validation.

**Feldmeyer Dirk:** Conceptualization, Writing – review & editing, Validation, Supervision, Resources, Project administration, Methodology, Funding acquisition.

**Koch Henner:** Writing – original draft, Writing – review & editing, Validation, Supervision, Resources, Project administration, Methodology, Funding acquisition, Conceptualization

