## Supplementary material for "Transport-Related Effects on Intrinsic and Synaptic Properties of Human Cortical Neurons: A Comparative Study": Suplementary data

### Supplementary Data

**Extended Figures: 2**

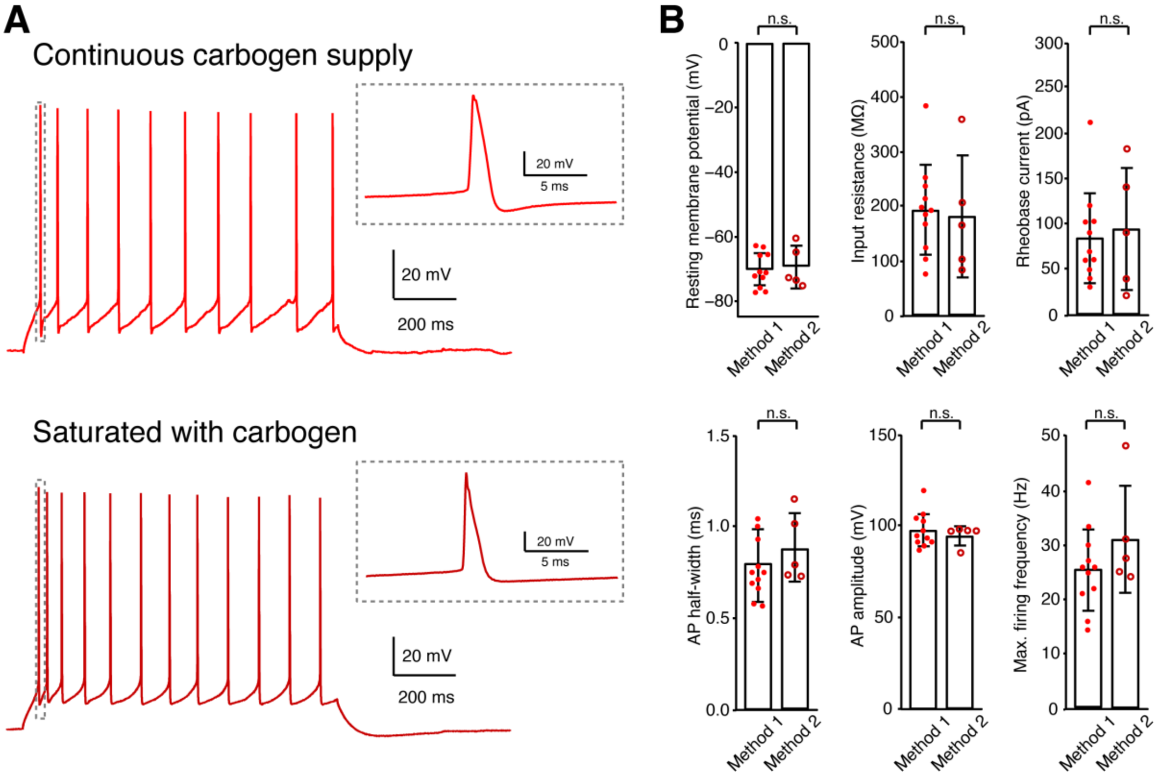


**Fig. 1-1. Comparison of electrophysiological properties of human cortical L2/3 pyramidal cells recorded in slices transported with continuous carbogen supply and saturated with carbogen.** (A) Representative firing patterns of human cortical L2/3 pyramidal cells recorded in slices transported with continuous carbogen supply (method 1, upper) and saturated with carbogen (method 2, lower). Insets, zoom-in of 1st APs. (B) Comparison of electrophysiological properties of pyramidal cells recorded in slices transported with method 1 (n=11) and method 2 (n=5). n.s. $p \geq 0.05$ for Wilcoxon Mann-Whitney U-test. No significant differences in intrinsic properties (e.g. resting membrane property, AP half-width and amplitude etc) were found between L2/3 pyramidal cells recorded in slices transported with two methods.

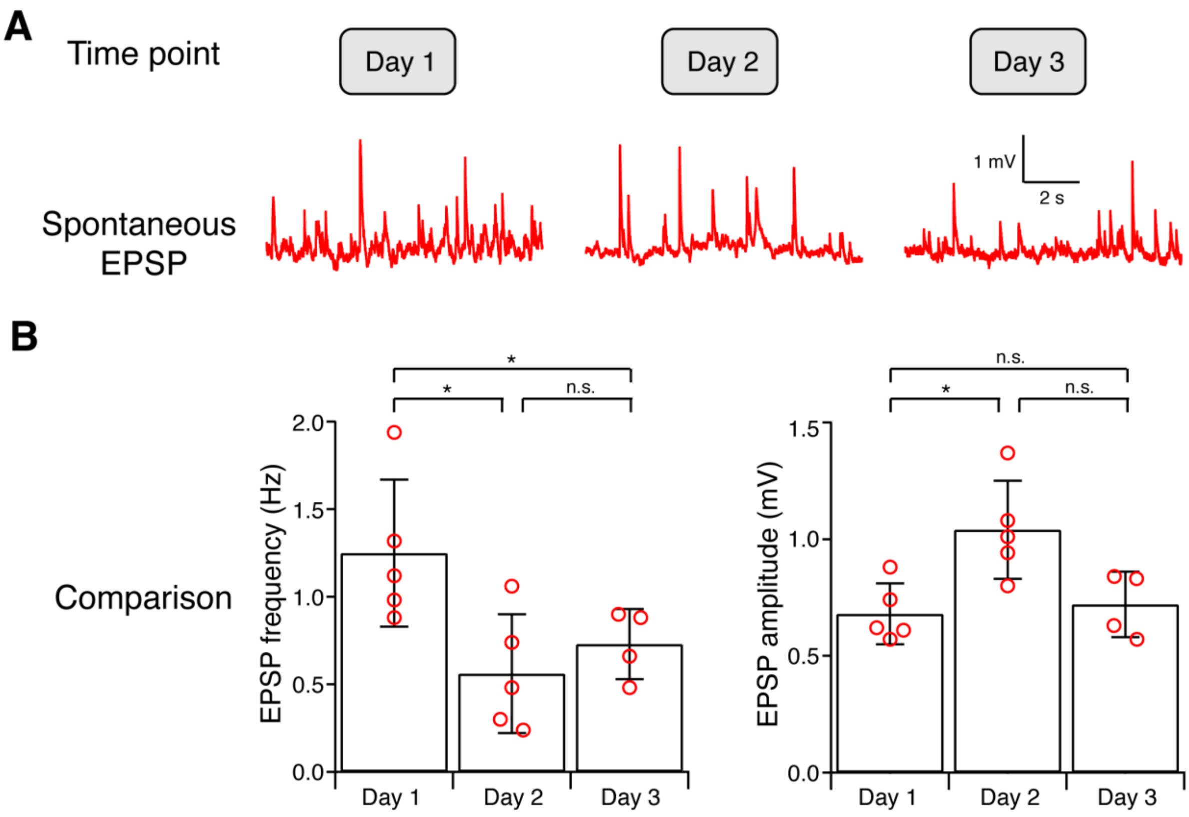


**Fig. 2-1. Changes of spontaneous EPSP frequency and amplitude for Off-site recorded human cortical neurons maintained in ex vivo for a prolonged (~ 60 hrs) time.** (A) Representative sEPSP recordings from cortical neurons maintained in ex vivo at three time points after the slice preparation. (B) Comparison of sEPSP frequecy and amplitude measured at three time points after the slice preparation. The sEPSP frequency significantly decreased at recordings performed at Day 2 and Day 3 compared to Day 1. The sEPSP amplitude increased in recordings obtained from Day 1 to Day2 and then decreased from Day 2 to Day 3; n=5 for Day 1, n=5 for Day2 and n=4 for Day 3. n.s. $p \geq 0.05$, * $p < 0.05$ for Wilcoxon-Mann-Whitney two-sample rank test.
